# Efficient sorting of single-unit activity from midbrain cells using KiloSort is as accurate as manual sorting

**DOI:** 10.1101/303479

**Authors:** Madeleine Allen, Tara Chowdhury, Meredyth Wegener, Bita Moghaddam

## Abstract

Extracting single-unit activity from *in vivo* extracellular neural electrophysiology data requires sorting spikes from background noise and action potentials from multiple cells in order to identify the activity of individual neurons. Typically this has been achieved by algorithms that employ principal component analyses followed by manual allocation of spikes to individual clusters based on visual inspection of the waveform shape. This method of manual sorting can give varying results between human operators and is highly time-consuming, especially in recordings with higher levels of background noise. To address these problems, automatic sorting algorithms have begun to gain popularity as viable methods for sorting electrophysiological data, although little is known about the use of these algorithms with neural data from midbrain recordings. KiloSort is a relatively new algorithm that automatically clusters raw data which can then be manually curated. In this report, we compare results of manually-sorted and KiloSort-processed recordings from the ventral tegmental area (VTA) and substantia nigra pars compacta (SNc). Sorting with KiloSort required substantially less time to complete, while yielding comparable and consistent results. We conclude that the use of KiloSort to identify single units from multi-channel recording in the VTA and SNc is accurate and efficient.

## Introduction

A necessary step in the analysis of neural electrophysiology data is the isolation of individual unit activity by the process of spike-sorting: grouping spikes from purported “single-units” while eliminating background noise and activity from other neurons (Harris et al., 2016). This has typically been done using manual user interfaces (Offline Sorter, Plexon; Xclust, M.A. Wilson; MClust, A.D. Redish), where operators must manually evaluate principal component analyses (PCAs) and individual waveform shapes in order to sort out the spikes of distinct units. Manual sorting presents two major challenges: it is time-consuming and subjective. Sorting spikes from one recording session can take several hours or more especially if it is a particularly noisy recording. The time it takes to manually sort has become less practical as high-density electrode probes with hundreds of channels are becoming more common (Berenyi et al., 2014; Du et al., 2011; Jun et al., 2017). Additionally, human intervention in spike sorting inherently causes problems of reproducibility and subjectivity. Manual sorting can yield error rates of up to 30% (Harris et al., 2000; Wood et al., 2004).

In an effort to increase efficiency and repeatability of sorting, automatic sorting algorithms have been gaining popularity despite skepticism within the field of neural electrophysiology (Harris et al., 2016). However, there is little published data showing the efficacy of automatic sorting algorithms with midbrain cells. We employed the sorting algorithm KiloSort to sort our data taken from electrodes placed in the ventral tegmental area (VTA) and substantia nigra pars compacta (SNc) and compared these results to data manually sorted by three highly-trained investigators.

## Methods

KiloSort is an open-source software that automatically clusters raw data and then functions with a manual user interface software for manual curation of the clusters. The algorithm runs through MATLAB, utilizing template matching during spike detection and spike clustering (Pachitariu et al., 2016). KiloSort utilizes a Graphical Processing Unit (GPU) to decrease run time (Pachitariu et al., 2016).

After running through KiloSort’s algorithm in MATLAB, the automatically clustered data is displayed in a Graphical User Interface (GUI) where an operator must manually curate the clusters (Figure 1). This involves merging and splitting the clusters as well as marking them as “good,” “MUA” (multi-unit activity), and “noise.” This classification simplifies data extraction after manual curation.

**Figure 1.**
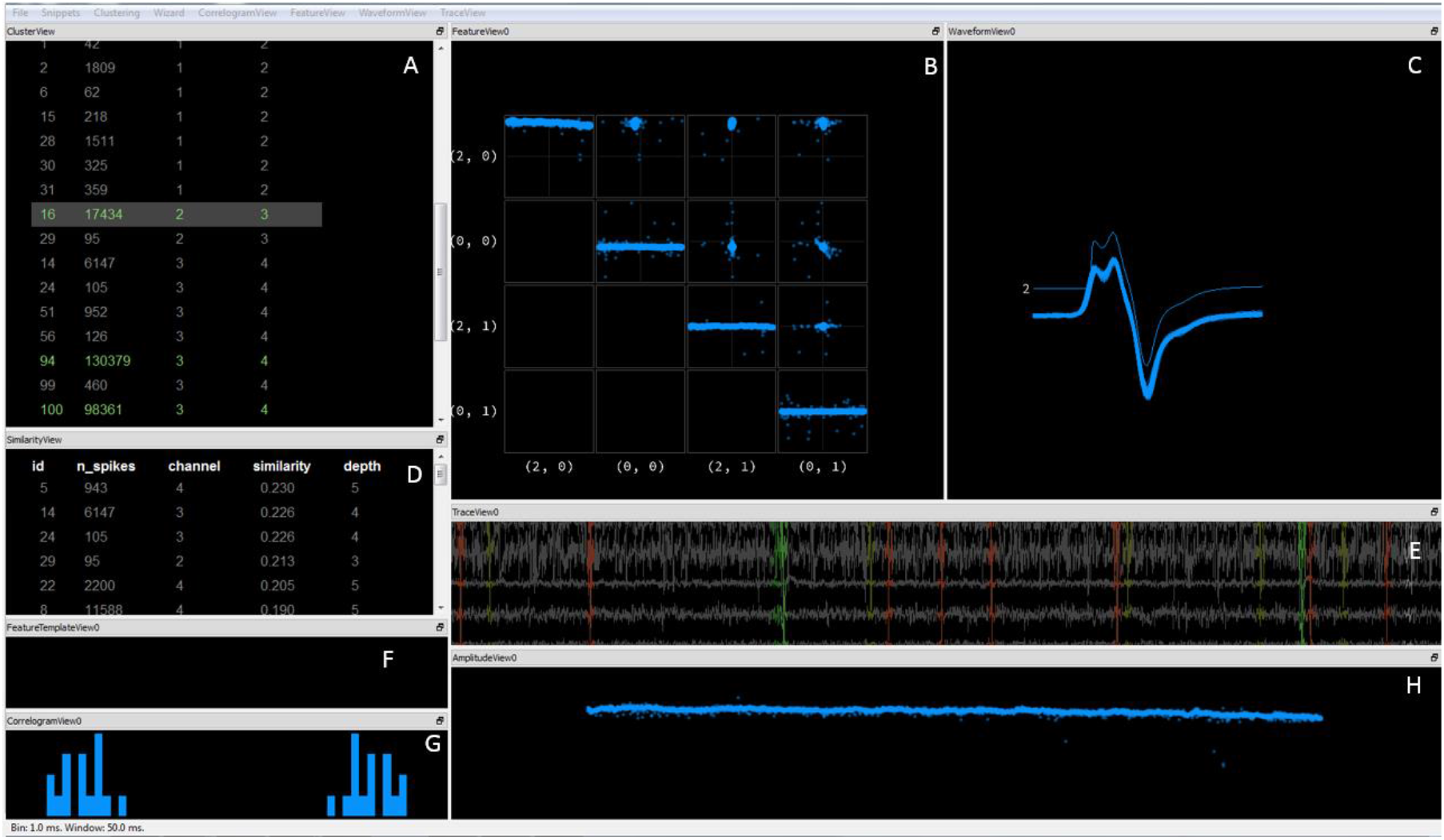
Template GUI used for manual curation in KiloSort. (A) Cluster View displays a list of all detected clusters and their ID number, number of spikes, channel it was derived from, and channel depth. Classifying a cluster as “good” will turn its label green (as shown), and classifying it as “MUA” or “noise” will turn it gray. Unclassified clusters display as white. (B) Feature View plots principal components of selected cluster. (C) Waveform View shows a plot of a subset of events of a selected cluster(s). (D) Similarity View is similar to Cluster View—it also displays all detected clusters and associated information, but in order of similarity to selected cluster in Cluster View. (E) Trace View displays plots of raw traces for each channel. Detected spikes are highlighted in different colors; associated events are highlighted in blue when a specific cluster is selected in Cluster View. (F) Feature Template View. (G) Correlogram View displays auto-correlograms of a selected cluster, and cross-correlograms of a pair of selected clusters. (H) Amplitude View plots template scaling for each event of a selected cluster against time (Lenzi & Steinmetz, 2017).

### Integration with Plexon data

Plexon’s Omniplex and MAP Offline SDK Bundle (Plexon) were used in MATLAB to convert Plexon data to raw binary files for KiloSort. Scripts from this package and from Kilosort were edited to import Plexon data into Kilosort. Custom MATLAB scripts were written to extract data from Kilosort. All of these scripts are freely available at github.com/madeleinea/KilosortPLX.

### Parameters

Kilosort includes a script (StandardConfig_MOVEME.m) with default parameters that should be changed based on data collection protocols. In sorting our data, almost all of the default parameters remained the same as previously set by Kilosort’s programmers, except for some parameters related to our system (firing rate, GPU availability). Otherwise, there were two major parameters that needed to be adjusted to achieve adequate spike sorting outcomes for our midbrain recordings (Table 1).

**Table 1.**
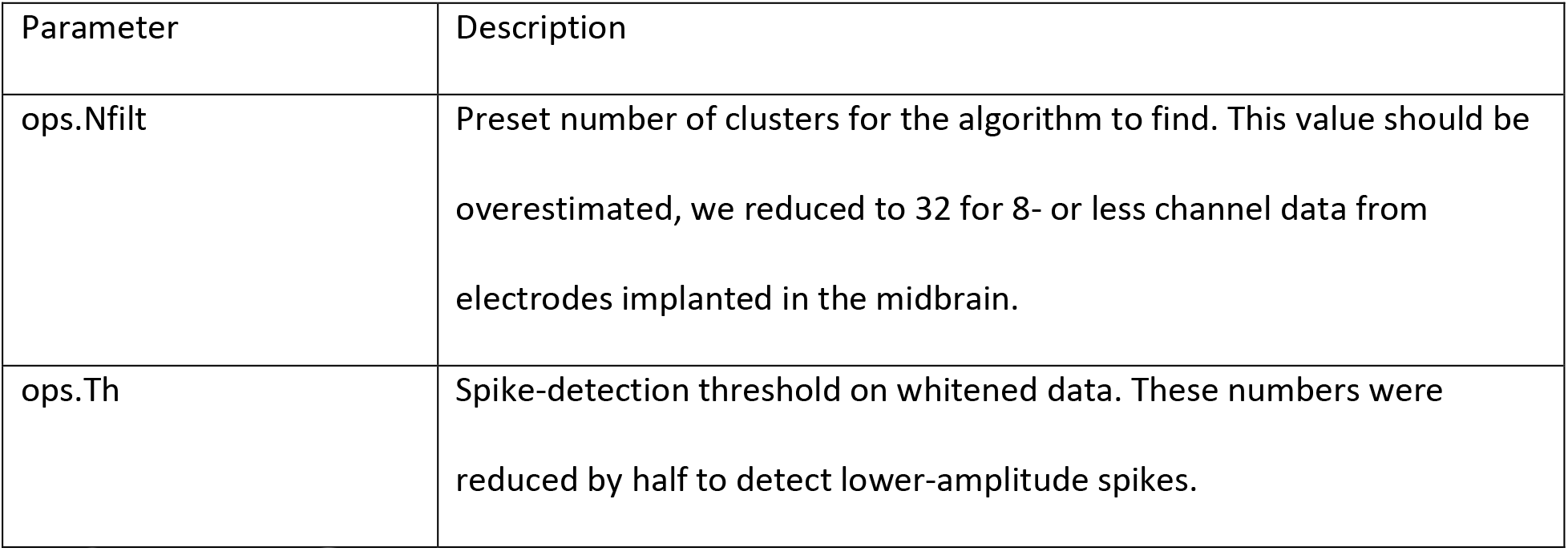
Parameters adjusted in KiloSort to improve accuracy. Two parameters were adjusted to achieve adequate results with recordings from the VTA and SNc.

### Technical Equipment

All data analysis and calculations were performed in MATLAB 2016b (Mathworks, Natick, MA) and Microsoft Excel (Microsoft Corp.) on a Dell Precision T1700 running 64-bit Windows 7 Enterprise with a 3.60 GHz Intel Core i7-4790 CPU, 32 GB of RAM, and a GeForce GTX 1050 Ti GPU (NVIDIA).

A GPU can be used with KiloSort by acquiring compatible versions of MATLAB, computer operating system, CUDA, Visual Studio (compiler), and a CUDA-enabled GPU. Visual Studio Professional 2013, MATLAB 2016b, CUDA 7.5, Windows 7, and GeForce GTX 1050 Ti were used in these analyses. A script included in the KiloSort script repository, mexGPUall.m, was then run to configure settings with the GPU.

### Data Analysis

Files from five 60-90 minute recordings with 6-8 channels were sorted manually in KiloSort, resulting in 26 units detected by both methods and 5 additional units detected only by KiloSort. All manually sorted data were sorted using Offline Sorter (Plexon) by one of three operators. All data sorted in KiloSort were curated by one operator in the GUI.

The percent difference in peaks and valleys per unit was used as a way to measure the difference in waveform features. This was found with the following equation:

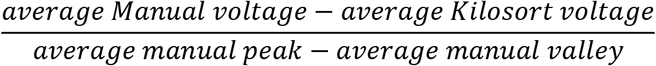

This equation finds the difference between the peak or valley voltages found with each sorting method, divided by the voltage range of the unit as defined by the difference between the average manual peak and valley voltages.

Two-tailed equal-variance Student’s t-tests were performed to compare averages (the number of units found per file and the number of waveforms found per unit in manually-sorted and KiloSort-processed data).

Twenty-nine 85-105 minute 8-channel recordings were run through KiloSort, and runtimes were recorded. Average times were calculated in Excel 2013.

### Data Acquisition

8-channel datasets were collected from male and female Long Evans and Sprague Dawley rats. Single-unit activity was recorded from behaving rats using custom microelectrode arrays of eight polyamide-insulated tungsten wires (50 μm). Signals were amplified using a multichannel amplifier (Plexon), and spikes were bandpass filtered between 220 Hz and 6 kHz, amplified 500-1000x gain, digitized at 40 kHz, and high-pass filtered at 300-8000 Hz.

## Results

Runtimes through KiloSort on our system averaged at 22.52 minutes per recording (SE = 2.42) for 29 8-channel recordings of 85-105 minutes. Manual curation of KiloSort-processed data typically takes 5 to 45 minutes. This is in contrast to manual sorting, which can take several hours to sort the same file.

Comparable results were obtained between data from manual sorting and data run through KiloSort. All of the same units as defined by visual waveform shape, number of events detected, and average peak and valley voltages were detected in KiloSort as in manual sorting (Fig. 2a). Similar numbers of events per mutual unit (a unit detected through both manual sorting and KiloSort) were detected in KiloSort and through manually sorting (KiloSort: *mean ± SE* = 34063.96 ± 11690.52; Offline Sorter: *mean ± SE* = 34116.54 ± 11844.71; *t (DF)* = 0.05, *p* = 0.997) (Fig. 2b). In addition, similar numbers of units were found per recording in both manners of sorting (KiloSort: *mean ± SE* = 6.2± 1.39; Offline Sorter: *mean ± SE* = 5.2 ± 1.31; *t (DF)* = 0.05, *p* = 0.616) (Fig. 2c). On average, there were small percentage differences in peaks and valleys between manually-sorted and KiloSort-processed results (Peaks: *mean ± SE* = 2.90% ± 1.16%; Valleys: *mean ± SE* = 1.62% ± 0.54%; *t (DF)* = 0.05, *p* = 0.997) (Fig. 2d). The global firing rates (firing rate across the entire recording) found through manual sorting and through Kilosort were similar and constant (Adjusted r-squared = 0.9985) (Fig. 2e).

**Figure 2.**
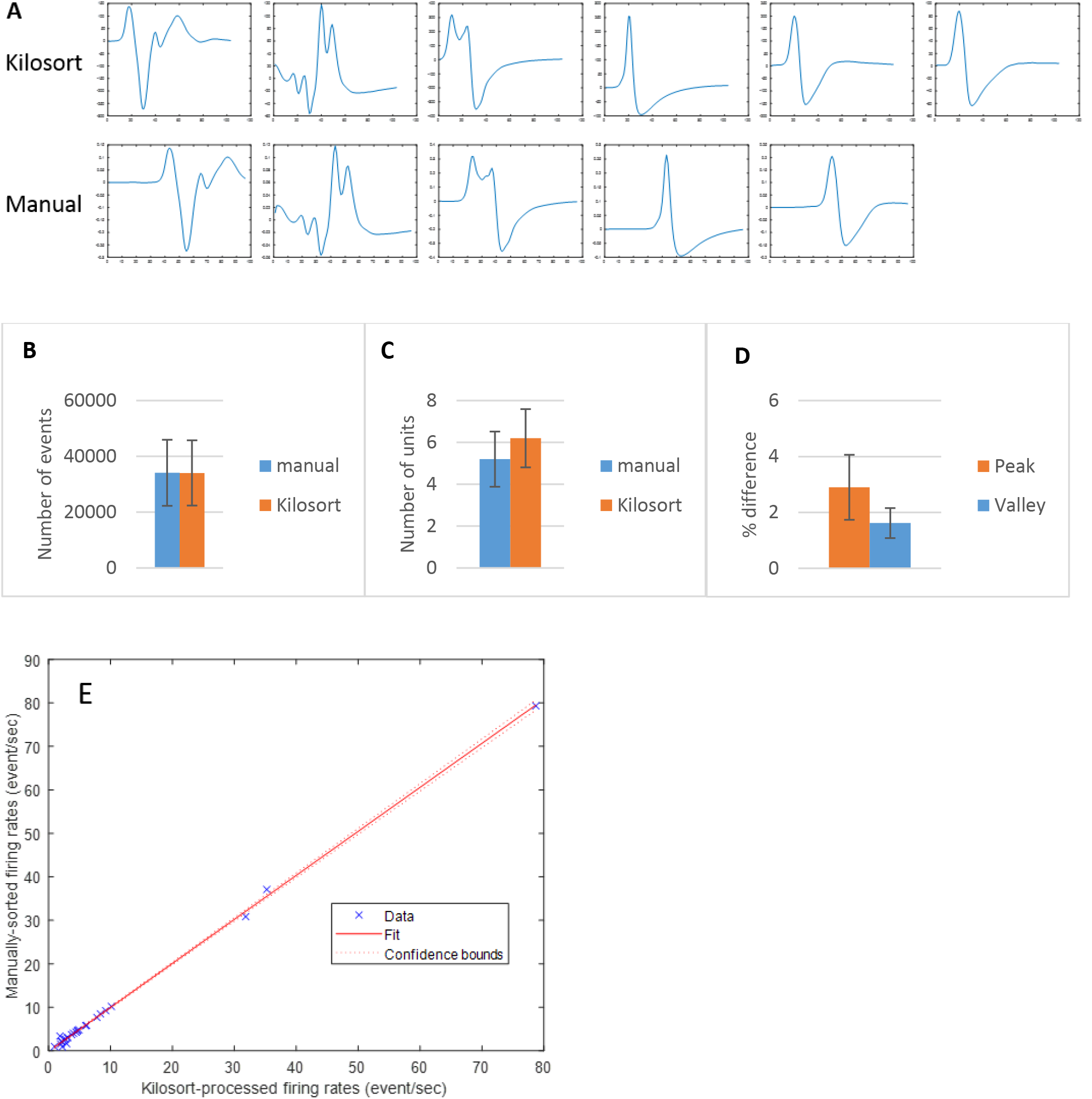
Similar results found in KiloSort and manual sorting. (A) Matched units detected through KiloSort and manual sorting from one recording. (B) Average number of waveform events detected per unit (of note, the bars appear to be identical because they are very close in value, supporting the notion that the two methods work similarly). (C) Average number of units detected per recording. (D) Average percent difference in valley and peak voltage between KiloSort and manual sorting. (E) Global firing rates per unit found in manual sorting and Kilosort. A linear fit within 95% confidence intervals was applied and plotted. The regression line equation is: f(x) = 1.01x - 0.16).

Several challenges of the GUI include the following: (1) only a subset of the waveforms are displayed, (2) merging/splitting of clusters can only be done in Feature View (Fig. 1b), and (3) there is no way to align waveforms in the GUI. The option to align waveforms can be important for the final waveform shape of a unit (Fig. 4). The GUI only displays a subset of a cluster’s waveforms in Waveform View (Fig. 1c), often hiding events that should be eliminated from a cluster. Figure 1c derives from the same data as in Figure 3, but displays only a portion of the events. In Figure 3a, obvious noise or alternate cluster events are present, but in the GUI they are invisible (Fig. 1c).

**Figure 3.**
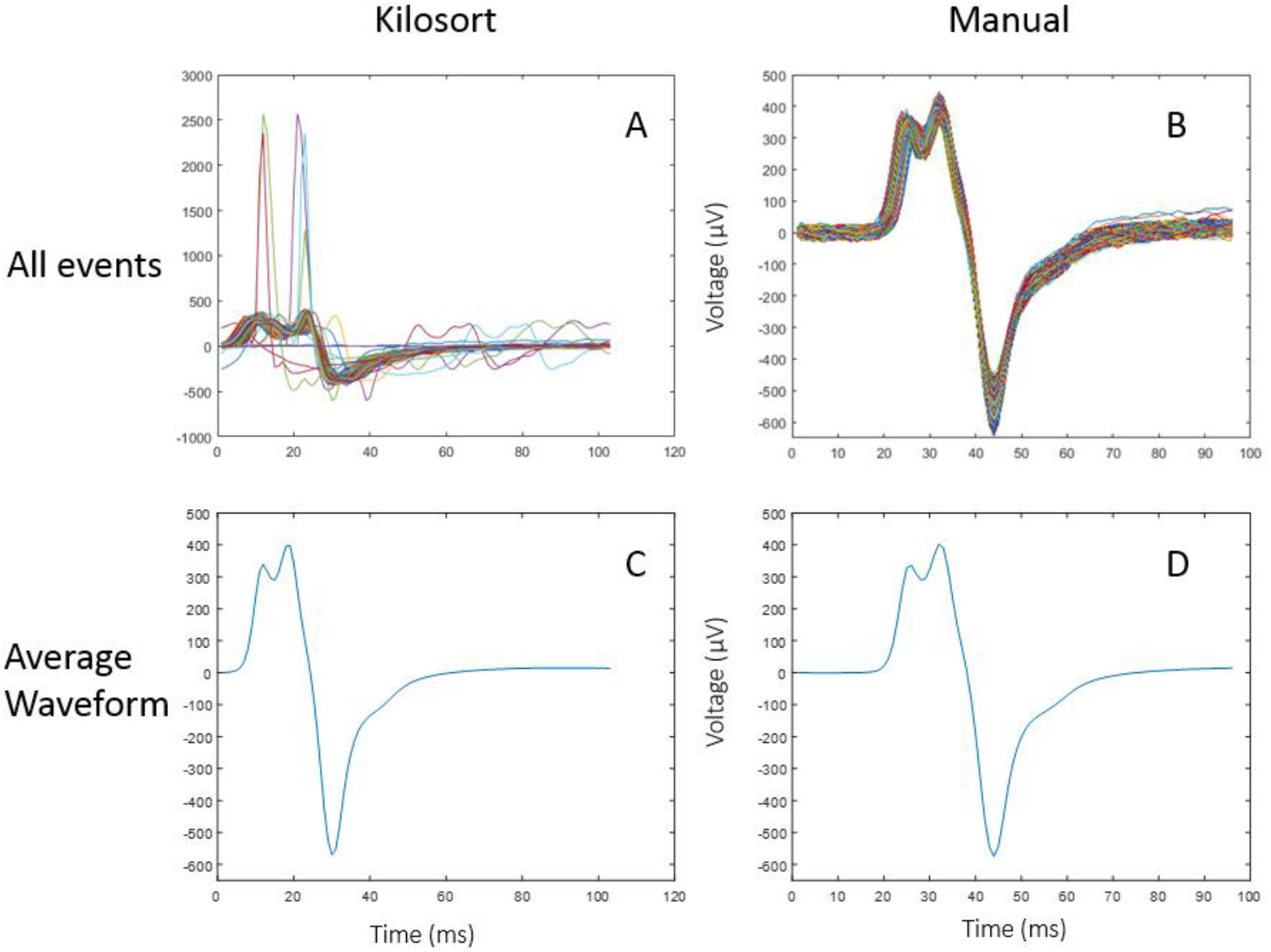
Resulting unit from KiloSort and manual sorting. (A) Plot of all events from unit detected in KiloSort. Several events are present that clearly should not be included in the final unit. (B) Plot of all events from unit detected in manual sorting. (C) Plot of average waveform event of unit found in KiloSort. (D) Plot of average waveform event of unit found in manual sorting.

**Figure 4.**
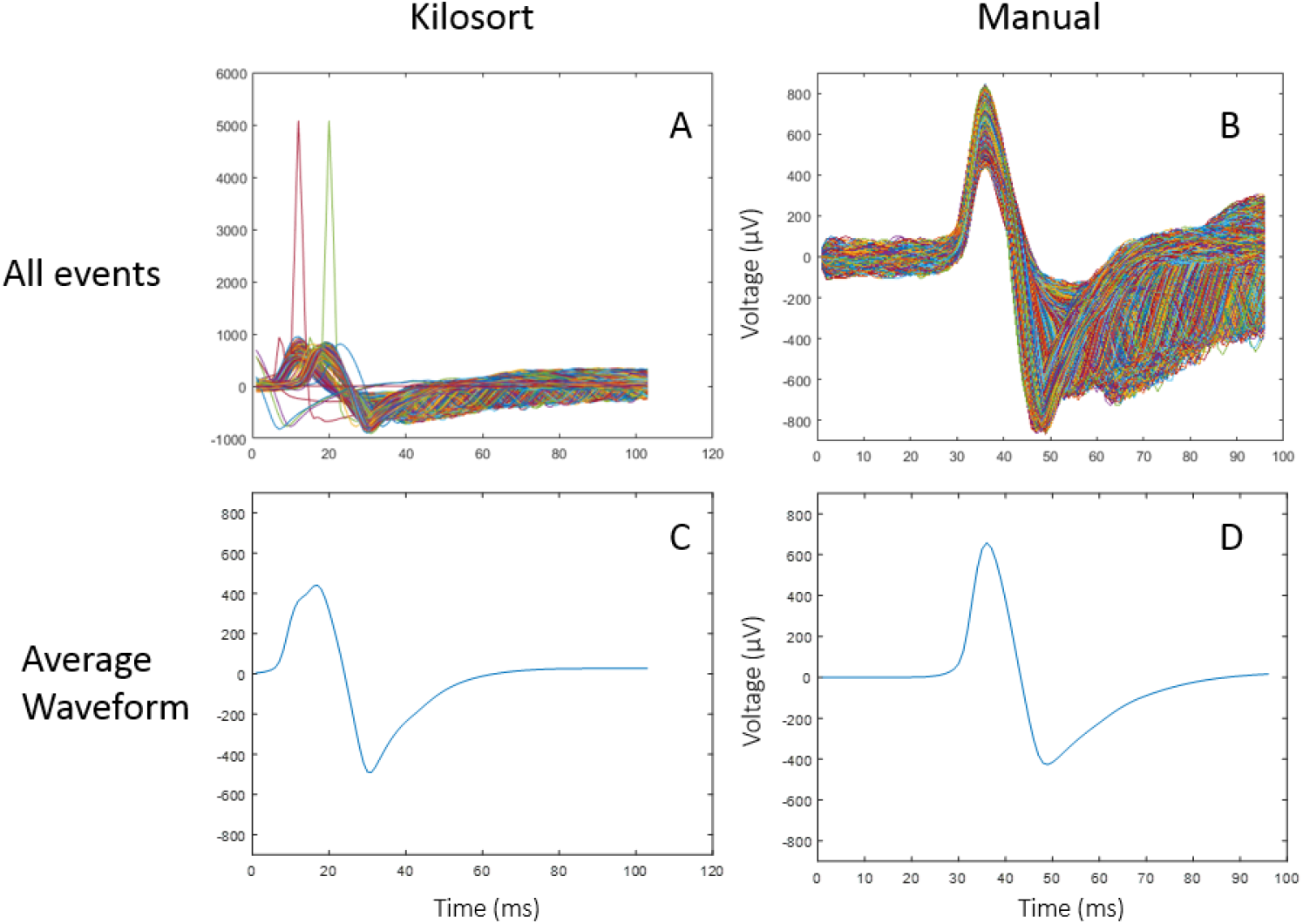
All events and average waveform plots of the same unit found in KiloSort and through manual sorting. (A) Plot of all events from unit detected in KiloSort. Some waveforms are out of alignment. (B) Plot of all events from unit detected in manual sorting. (C) Plot of average waveform event of unit found in KiloSort. (D) Plot of average waveform event of unit found in manual sorting.

The option to split clusters during manual curation is possible in the GUI, however this can only be done in Feature View (Fig. 2b). The operator can draw shapes around events (plotted as dots in Feature View) to split clusters. Clusters can also be merged in the GUI.

Additionally, there is no feature available to align waveforms in the GUI, although on occasion a substantial portion of events will be less than 0.25 ms out of alignment from the rest of the events of the same unit. The percent differences in peak and valley voltages between manual sorting and KiloSort for the misaligned unit in Figure 4 are 23.31% and 6.81%, respectively. One unit out of 26 units in this analysis (3.85%) had a notable number of misaligned events.

## Discussion

### Efficiency

Sorting using KiloSort takes considerably less time than manual sorting. While running multiple recordings through KiloSort could take hours, this is inactive time and the algorithm can simply be programmed to automatically run through files overnight. The active time spent manually curating clusters is largely reduced in comparison to manual sorting.

### Accuracy

The differences between results derived from manual sorting and KiloSort are negligible. The same units are found in KiloSort as in manual sorting (Fig. 2a), similar numbers of events are found per unit (Fig. 2b), and important features of the waveforms shapes, such as peak and valley voltages, are similar (Fig. 2d).

KiloSort is more reliable than manual sorting because it reduces the need for human input and subjectivity, increasing consistency and repeatability. KiloSort will compute the same clusters every time it is run for the same file, whereas manual operators’ sorting results vary between occasions and across operators (Wood et al., 2004). Although manual curation is required with use of KiloSort, human interference is minimal compared to that of fully-manual sorting. The clustering provided by the KiloSort algorithm generally only requires the elimination of a few dozen events from a cluster or the merging with other clusters. The vast majority of the analysis included in sorting is computed by KiloSort.

Although Kilosort was designed to sort high-density electrode arrays, we found that it is efficient and effective with our 8-channel recordings. The automatic aspect and the configuration of the GUI optimize Kilosort for high-density electrode arrays—voltage traces from a unit are not only plotted in the channel containing the strongest signal, but all channels (1c). In this way, users can easily recognize if a voltage change was detected in multiple channels and identify the channel with the strongest signal.

### Challenges with manual curation

One problem of the GUI is that it yields “messier” units than those found in manual sorting, in the sense that several events are included in the final unit that should clearly have been eliminated (Fig. 3a). This occurs because (1) operators are often unable to see events that should be eliminated from a cluster, and (2) if events that should be eliminated are identified in Waveform View, the operator must then use Feature View to identify the associated points to eliminate these events from the cluster. This can be challenging, as there is no way to definitively link events plotted in Waveform View to those plotted in Feature View. A helpful next step for the GUI would be to enable splitting clusters in Waveform View rather than having this ability limited to Feature View.

The “messy” units from KiloSort, however, may not meaningfully affect further data analysis. We observed only a few stray events that do not affect the average waveform shape (Fig. 3c), and only slightly affect the timestamps and firing rate of a unit, which are arguably the most important data to extract from raw recordings.

About 3.8% of the time, a large portion of events from a cluster found in KiloSort were misaligned with the rest of the events of that cluster. This was caused by some spikes being out of alignment by a very small amount (Fig. 4a). This does not considerably affect the timestamps or firing rate of the unit, but it can affect the average waveform shape (Fig. 4c). For the misaligned unit in Figures 4a and 4c, the differences in peak and valley voltages between KiloSort and manual sorting are 3- to 10-fold higher than the average differences in peak and valley voltage for all units in this analysis. This demonstrates that the issue of alignment can have effects on the average waveform shape of a misaligned unit. The average peak voltage is especially affected in this case, because the misalignment causes a spread in the timestamps of the peak voltages. In contrast, Offline Sorter allows the operator to align waveforms using the peak voltage of individual events, creating a common reference point.

## Conclusion

We show that use of KiloSort is appropriate and may be more consistent than manual sorting of electrophysiology data on cells of the VTA and SNc, many of which are putative dopamine neurons. Overall we observed that KiloSort consistently yields results similar to those found in manual sorting, but is much more readily reproducible and time-efficient. The only issue with accuracy that arose in this analysis is that of misaligned events. But this happens rarely, and does not have a significant effect on firing rate and individual waveform shape of the present dataset. This problem could also potentially be resolved in post-processing steps.

